# Inhibition of guanosine monophosphate synthetase (GMPS) blocks glutamine metabolism and prostate cancer growth *in vitro* and *in vivo*

**DOI:** 10.1101/2020.09.07.286997

**Authors:** Qian Wang, Yi F. Guan, Sarah E. Hancock, Kanu Wahi, Michelle van Geldermalsen, Blake K. Zhang, Angel Pang, Rajini Nagarajah, Blossom Mak, Lisa G. Horvath, Nigel Turner, Jeff Holst

**Affiliations:** Translational Cancer Metabolism Laboratory, School of Medical Sciences and Prince of Wales Clinical School, UNSW Sydney, Australia; Mitochondrial Bioenergetics Laboratory, Department of Pharmacology, School of Medical Sciences, UNSW Sydney, Sydney, Australia; Origins of Cancer Program, Centenary Institute, University of Sydney, Camperdown, New South Wales, Australia; Sydney Medical School, University of Sydney, Sydney, New South Wales, Australia; Chris O’Brien Lifehouse, Sydney, Australia; Garvan Institute of Medical Research, Sydney, Australia; University of NSW, Sydney, Australia; University of Sydney, Sydney, Australia

## Abstract

Cancer cells increase their uptake of nutrients and metabolize them to provide the necessary building blocks for new cancer cells. Glutamine is a critical nutrient in cancer, however its contribution to purine metabolism in prostate cancer has not previously been determined. Guanosine monophosphate synthetase (GMPS) acts in the *de novo* purine biosynthesis pathway, utilizing a glutamine amide to synthesize the guanine nucleotide and replenish the purine pool in proliferative cancer cells. This study demonstrates that GMPS mRNA expression correlates with Gleason score in prostate cancer samples, while high GMPS expression was associated with decreased rates of overall and disease/progression-free survival. Pharmacological inhibition or knockdown of GMPS significantly decreased cell growth in both LNCaP and PC-3 prostate cancer cells. GMPS knockdown was rescued by addition of extracellular guanosine to the media, suggesting a direct effect on nucleotide synthesis. We utilized ^15^N-(amide)-glutamine and U-^13^C_5_-glutamine metabolomics to dissect the pathways involved, and intriguingly, despite similar growth inhibition by GMPS knockdown, we show unique metabolic effects across each cell line. PC-3 cells showed a build-up of purine precursors, as well as activation of purine salvage pathways highlighted by significant increases in guanine, adenosine, inosine and cytosine. Both cell lines exhibited increased levels of pyrimidines and prioritized TCA cycle in distinct ways to produce increased aspartate, another important purine precursor. Using a PC-3 xenograft mouse model, tumor growth was also significantly decreased after GMPS knockdown. These data further highlight the importance of glutamine metabolism for prostate cancer cell growth and provide support for GMPS as a new therapeutic target in prostate cancer.

## Introduction

Despite an understanding of cancer metabolism that spans almost 100 years since the Warburg effect was first described, only a few therapeutics targeting this key vulnerability have emerged. Cancer cells greatly increase their uptake of nutrients (glucose, amino acids and lipids), and metabolize them to provide the necessary building blocks (membranes, DNA, RNA, protein etc.) for new cancer cells. It has previously been shown that extracellular amino acids make up by far the majority of the carbon sources used by cancer cells for cell division, highlighting amino acid uptake and metabolism as a viable therapeutic target (Hosios et al., 2016).

The availability of glutamine and other amino acids is critical to sustain proliferation in cancer cells by providing the building blocks for protein, nucleotide and glutathione synthesis (Christensen, 1990; Kovacevic and McGivan, 1983). We have previously shown that expression of the glutamine transporter ASCT2 is increased in melanoma, prostate cancer, breast cancer and endometrial carcinoma in order to facilitate the increased glutamine demand required for tumor growth (Marshall et al., 2017; van Geldermalsen et al., 2016; Wang et al., 2014; Wang et al., 2015). Selective targeting of the ASCT2 transporter by antisense/shRNA inhibited glutamine uptake, resulting in decreased cellular viability in cancer cells (Marshall et al., 2017; van Geldermalsen et al., 2016; Wang et al., 2014; Wang et al., 2015). In prostate cancer, ASCT2 expression is regulated by androgen receptor transcription, as well as adaptive responses to nutrient stress mediated by ATF4 (Wang et al., 2013); and ASCT2 is also known to be regulated by MYC (Wise et al., 2008). These transcriptional pathways sustain high levels of ASCT2 expression and ASCT2-mediated glutamine uptake in prostate cancer, fueling their glutamine addiction.

Glutamine can be utilized for the TCA cycle through its conversion to glutamate (via glutaminase; GLS) and then to α-ketoglutarate (α-KG). This vulnerability has been targeted through glutaminase inhibitors in a range of cancer subsets such as triple-negative breast cancer (Gross et al., 2014), and more recently in a metastatic subline of PC-3 cells, PC3-M (Zacharias et al., 2017). Glutamine dependency has also been shown to drive neuroendocrine differentiation in prostate cancer, with ASCT2 inhibition decreasing tumor growth (Mishra et al., 2018). Along with this important role for glutamine carbons in prostate cancer, glutamine also acts as a nitrogen donor for *de novo* purine and pyrimidine biosynthesis. The purine *de novo* pathways are sequentially orchestrated by six enzymes, which are co-localized to form the purinosome, to catalyze the conversion of phosphoribosylpyrophosphate (PRPP) to inosine 5’-monophosphate (IMP)(An et al., 2008). For proliferating cells, which require large amount of purine nucleotides for DNA and RNA synthesis, the *de novo* biosynthetic pathway is important to replenish the purine pool. In addition to the *de novo* biosynthetic pathway, purine nucleotides can also be synthesized through the salvage pathway in mammalian cells. The salvage pathway produces purines by recycling the degraded bases through the activity of hypoxanthine-guanine phosphoribosyltransferase (HPRT1) and adenine phosphoribosyltransferase (APRT) (Stout and Caskey, 1985). HPRT1 recycles both hypoxanthine and guanine to generate inosine monophosphate (IMP) and guanine monophosphate (GMP).

Guanosine monophosphate synthetase (GMPS) is one of three glutamine amidotransferases involved in *de novo* purine biosynthesis and is responsible for the last step in the synthesis of the guanine nucleotide, GMP. It catalyzes the conversion of xanthine monophosphate (XMP) to GMP in the presence of glutamine and ATP. Human GMPS has 693 residues, containing two functional domains: the N-terminal glutaminase (GATase) domain and the C-terminal synthetase (ATPPase) domain (Tesmer et al., 1996; Welin et al., 2013). Glutamine binds to the GATase domain of GMPS and generates ammonia, which aminates XMP at the ATPPase domain to generate GMP (Welin et al., 2013). GMPS has been shown to be overexpressed in metastatic human melanoma cells, where GMPS inhibition suppresses melanoma cell invasion and tumorigenicity (Bianchi-Smiraglia et al., 2015), but has not previously been examined in prostate cancer.

In this study, we use clinical and cell line data to show that GMPS expression is upregulated and is functionally important for cancer cell growth, validating GMPS as a putative therapeutic target in prostate cancer. Furthermore, we uncover distinct glutamine metabolic pathways utilized by different prostate cancer cells, which may also be therapeutically targeted.

## Results

### GMPS expression in prostate cancer

To determine the expression of GMPS in prostate cancer, we first compared prostate cancer to matched normal samples in the TCGA dataset, showing a significant increase in GMPS mRNA expression (**Fig. 1A**; n=52). Similar significant increases were observed in the majority of prostate cancer datasets using Oncomine (**Table 1**, **Fig. 1B** **and** **1C**) (Arredouani et al., 2009; Grasso et al., 2012; Lapointe et al., 2004; Vanaja et al., 2003; Welsh et al., 2001). Interestingly, GMPS expression was also significantly increased in metastatic prostate cancer compared to primary prostate cancer (**Fig. 1B** **and** **1C**), and correlated with metastatic castration resistant prostate cancer cell cycle genes CDK1, CDC20 and UBE2C (**Fig. S1A**) (Wang et al., 2009; Wang et al., 2013). In the TCGA dataset, GMPS mRNA expression was significantly increased in patients with higher Gleason score cancers (**Fig. 1D**), with a significant decrease in overall and disease/progression-free survival in the ~15% of patients with GMPS alterations (Gain, amplification or mRNA Z-Score≥2; **Fig. 1E, 1F** **and** **Fig. S1B**). In addition, GMPS expression correlated with a number of additional enzymes involved in purine (PPAT, PFAS, GART and PAICS) and pyrimidine (CAD and CTPS1) biosynthesis and glutaminolysis (GLS; **Fig. S1C**), suggesting that many prostate cancer cells are primed with all the necessary machinery for glutamine usage for TCA and nucleotide synthesis. Taken together, these data suggest that high GMPS expression may play an important role in higher grade metastatic prostate cancer by regulating nucleotide synthesis.

**Table 1.** GMPS expression in Oncomine datasets (Cancer vs Normal)

| **Dataset** | **Fold Change** | **P-value** | **# PCa Samples** |
| --- | --- | --- | --- |
| Luo Prostate 2 | 1.871 | 2.01E-04 | 15 |
| Welsh Prostate | 1.598 | 5.33E-05 | 25 |
| Singh Prostate | 1.591 | 2.50E-02 | 52 |
| Tomlins Prostate | 1.588 | 3.00E-03 | 49 |
| Vanaja Prostate | 1.317 | 1.43E-04 | 32 |
| Grasso Prostate | 1.311 | 3.27E-09 | 59 |
| Arredouani Prostate | 1.227 | 1.60E-02 | 13 |
| Lapointe Prostate | 1.21 | 1.41E-05 | 71 |

**Figure 1.**
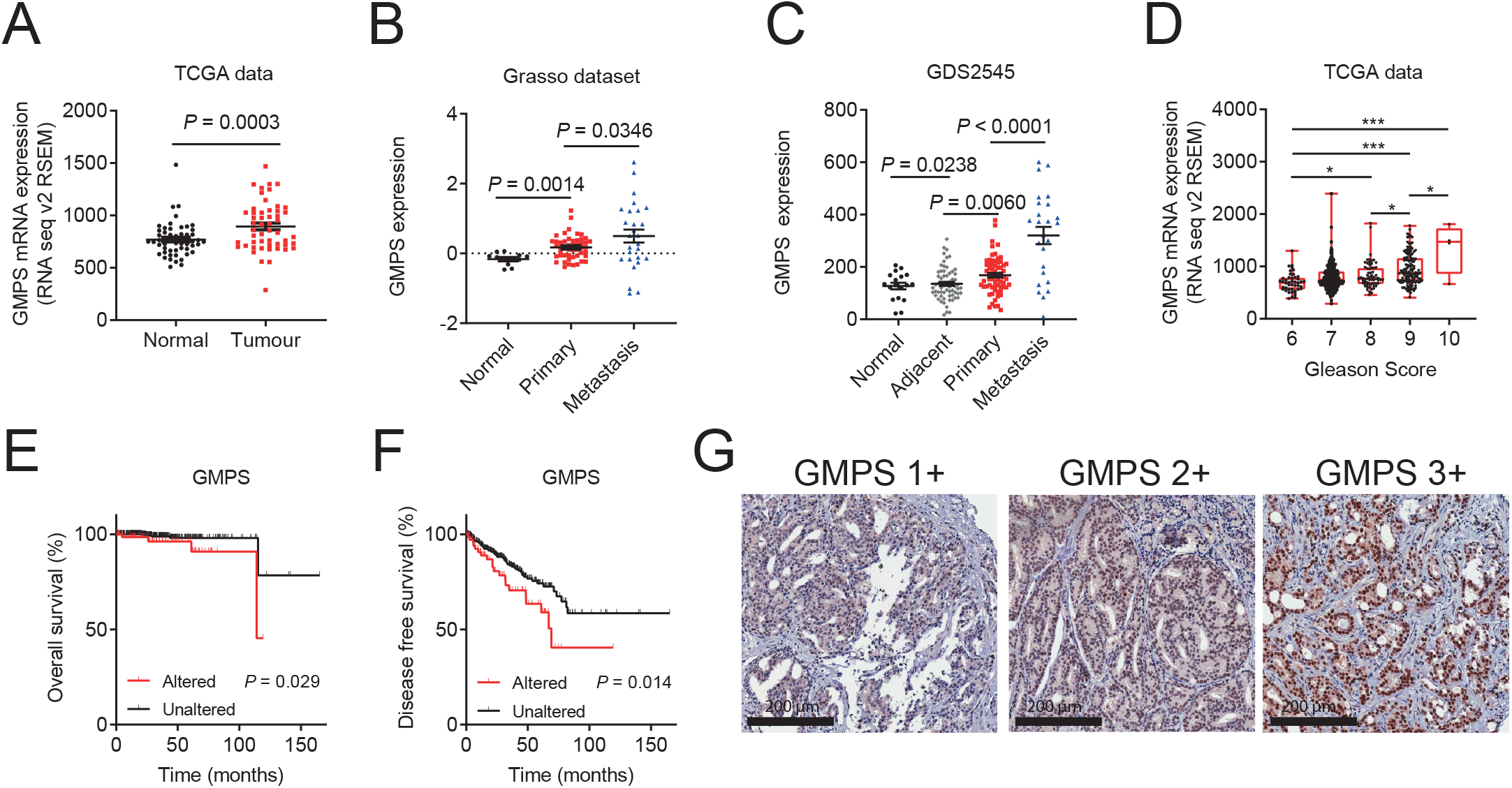
GMPS expression in prostate cancer patient cohorts. (A) GMPS mRNA expression in matched prostate cancer samples compared to adjacent normal prostate from the TCGA dataset. Data are mean ± SEM; paired t-test; n=52. (B) GMPS expression in normal prostate (n = 12), primary (n = 49) and metastatic prostate cancer (n = 27) from the Grasso dataset. Data are mean ± SEM; unpaired parametric t-test. (C) GDS2545 samples are from normal prostate (Normal), primary cancer adjacent normal (Adjacent), primary cancer (Primary) and androgen-ablation resistant metastatic prostate cancers (Metastatic). Data are mean ± SEM; unpaired parametric t-test. (D) GMPS mRNA expression was assessed for Gleason Score 6 (n=45), 7 (n=244), 8 (n=63), 9 (n=135) and 10 (n=4). Data are mean ± SEM; One-way ANOVA test; *, P < 0.05; **, P < 0.01; ***, P < 0.001. (E and F) Overall survival (E; altered=74, unaltered=417) and Disease-free survival (F; altered=73, unaltered=412) was assessed comparing patients with significant alterations in GMPS (Gain, amplification or mRNA expression Z-score≥2) to remaining patients using cBioPortal. (G) Representative images of GMPS protein expression by immunohistochemistry in prostate cancer patient samples. Scale bar is 200 μm.

In addition, we examined GMPS protein expression in a cohort of prostate cancer patient samples (n=63) using immunohistochemistry. GMPS protein was expressed in both the cytoplasm and nucleus of cancer cells (**Fig. 1G**). While there was a trend but no significant increase in the average IHC score for GMPS expression in different Gleason scores (**Fig. S1D**), there was a significant increase in the percentage of GMPS positive cells in high Gleason score ≥8 cancers (**Fig. S1E**). We also showed high protein expression of the glutamine transporter ASCT2 in this cohort (**Fig. S1F** **and** **S1G**), confirming these cancer cells have the transport capacity for high glutamine uptake.

### Inhibition of GMPS blocks cell growth without causing apoptosis

We next examined GMPS protein expression in LNCaP, PC-3 and DU145 human prostate cancer cell lines. GMPS was detected in all three cell lines, with highest expression in LNCaP and DU145 cells (**Fig. 2A**). To determine the function of GMPS in prostate cancer, we first used a nucleoside analog decoyinine (Deco) to inhibit GMPS activity. Decoyinine is a selective, reversible non-competitive GMPS inhibitor active between 50-200 μM (Nakamura and Lou, 1995). We initially confirmed that decoyinine was active in LNCaP cells at these concentrations, showing a half maximal inhibitory concentration (IC_50_) of 102.5 μM (**Fig. 2B**). To ensure maximal inhibition, 200 μM decoyinine was used in subsequent experiments. After treatment with decoyinine for three days, LNCaP, PC-3 and DU145 cells showed a significant reduction of cell growth compared to control group, similar to that achieved by complete glutamine starvation (**Fig. 2C, 2D** **and** **2E**), suggesting that GMPS activity is required for prostate cancer cell growth. To investigate if decoyinine induces apoptosis, we stained cells with Annexin V antibody and PI after two days treatment with decoyinine. LNCaP cells are sensitive to glutamine starvation and showed a significant increase in apoptosis, while only a mild increase was seen in PC-3 cells (**Fig. 2F** **and** **2G**). By contrast, decoyinine treatment did not induce apoptosis despite the substantial effects on cell growth (**Fig. 2F** **and** **2G**). These results suggested that GMPS is more important for prostate cancer cell proliferation.

**Figure 2.**
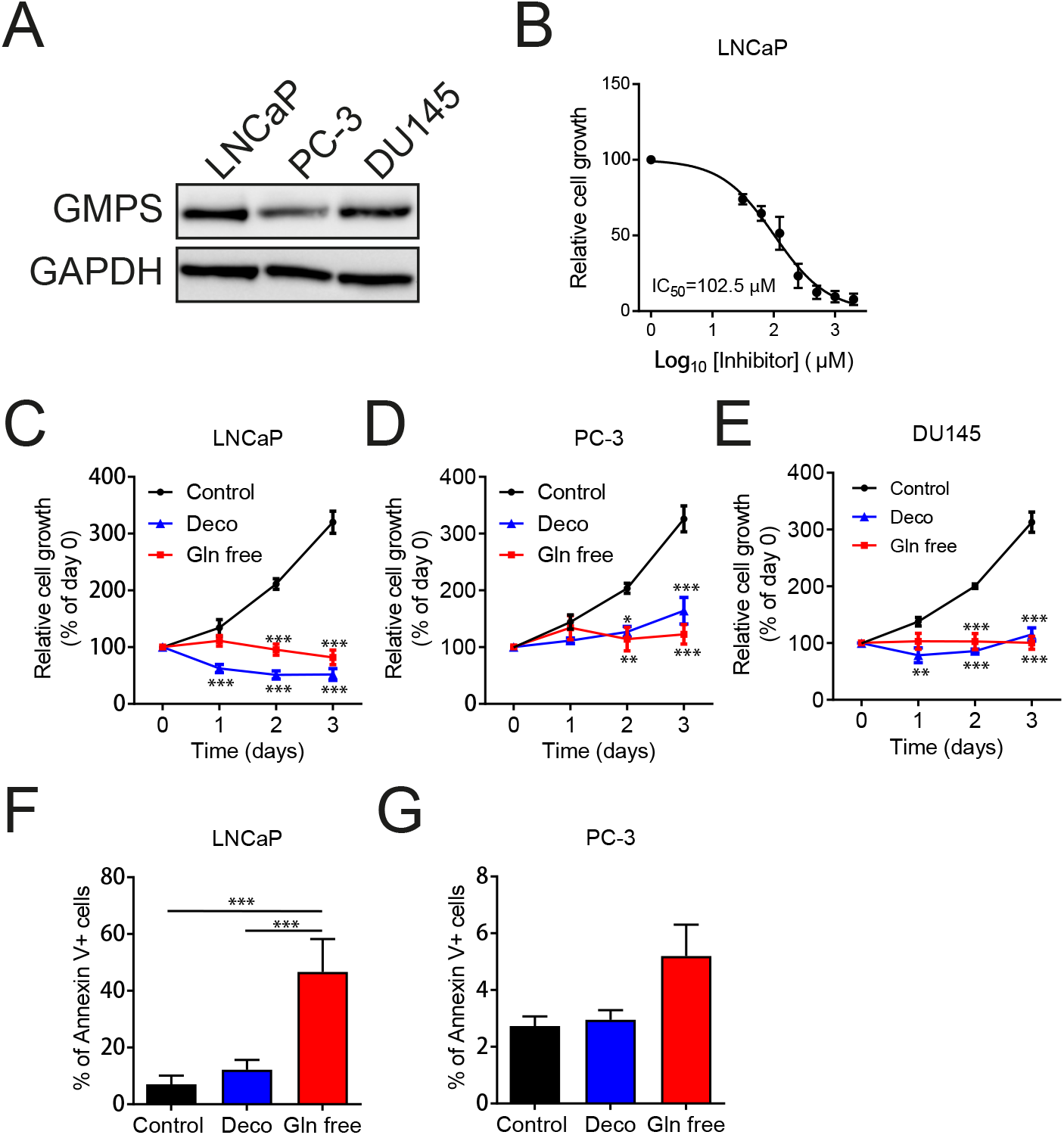
GMPS inhibitor decoyinine blocks prostate cancer cell growth. (A) GMPS protein was detected by western blotting in LNCaP, PC-3 and DU145 prostate cancer cell lines. GAPDH was used as the loading control. (B) Dose response of GMPS inhibitor decoyinine in LNCaP cells was assessed using an MTT assay. (C, D and E) Effects of decoyinine (Deco; 200 μM) or glutamine free media on prostate cancer cell growth in LNCaP (C, n=4), PC-3 (D, n=4) and DU145 (E, n=4) cell lines using an MTT assay. Data are mean ± SEM; Two-way ANOVA test. (F and G) LNCaP cells (F) or PC-3 cells (G) were cultured for 48 h in the presence or absence of decoyinine or in glutamine (Gln) free media. Cells were stained using an Annexin V antibody and examined by flow cytometry. Data are mean ± SEM. n=3. One-way ANOVA test. *, P < 0.05; **, P < 0.01; ***, P < 0.001.

### GMPS knockdown inhibits proliferation and can be rescued by the purine salvage pathway

To better understand the function of GMPS, we used shRNA targeting GMPS to decrease GMPS expression levels. Two different lentiviral shRNAs, shGMPS-41 and shGMPS-42 were used to transduce LNCaP and PC-3 cells, with both achieving effective GMPS protein knockdown (**Fig. 3A** **and** **3B**). We also examined the GMPS localization using immunofluorescence staining, with GMPS detected in both the cytoplasm and nucleus (**Fig. S2A**), consistent with our IHC staining in patient samples (**Fig. 1G**). After shRNA knockdown, GMPS levels were substantially decreased in both the cytoplasm and nucleus in shGMPS cells compared to shControl cells (**Fig. S2A**). GMPS knockdown significantly decreased viable cell numbers in LNCaP and PC-3 cells (**Fig. 3C** **and** **3D**), supporting the decoyinine data. To more directly determine the effects on proliferation, BrdU incorporation was also assessed. In LNCaP cells, shGMPS-41 and shGMPS-42 decreased cell proliferation to 60.7% and 54.3% of shControl cells, respectively, which was similar to Gln free media treated cells (50.4% of shControl cells; **Fig. 3E**). In PC-3 cells, shGMPS-41 and shGMPS-42 decreased cell proliferation to 76.9% and 70.1% of shControl cells, respectively (**Fig. 3F**). However, in PC-3 cells, Gln free media treated cells showed a more profound inhibition (25.1% of shControl cells; **Fig. 3F**). Similar to decoyinine, neither shGMPS-41 and shGMPS-42 were able to induce apoptosis in LNCaP or PC-3 cells (**Fig. S2B** **and** **S2C**), nor were there any significant changes in cleaved PARP (c-PARP; **Fig. S2D**) further suggesting that GMPS knockdown did not induce apoptosis in prostate cancer cells.

**Figure 3.**
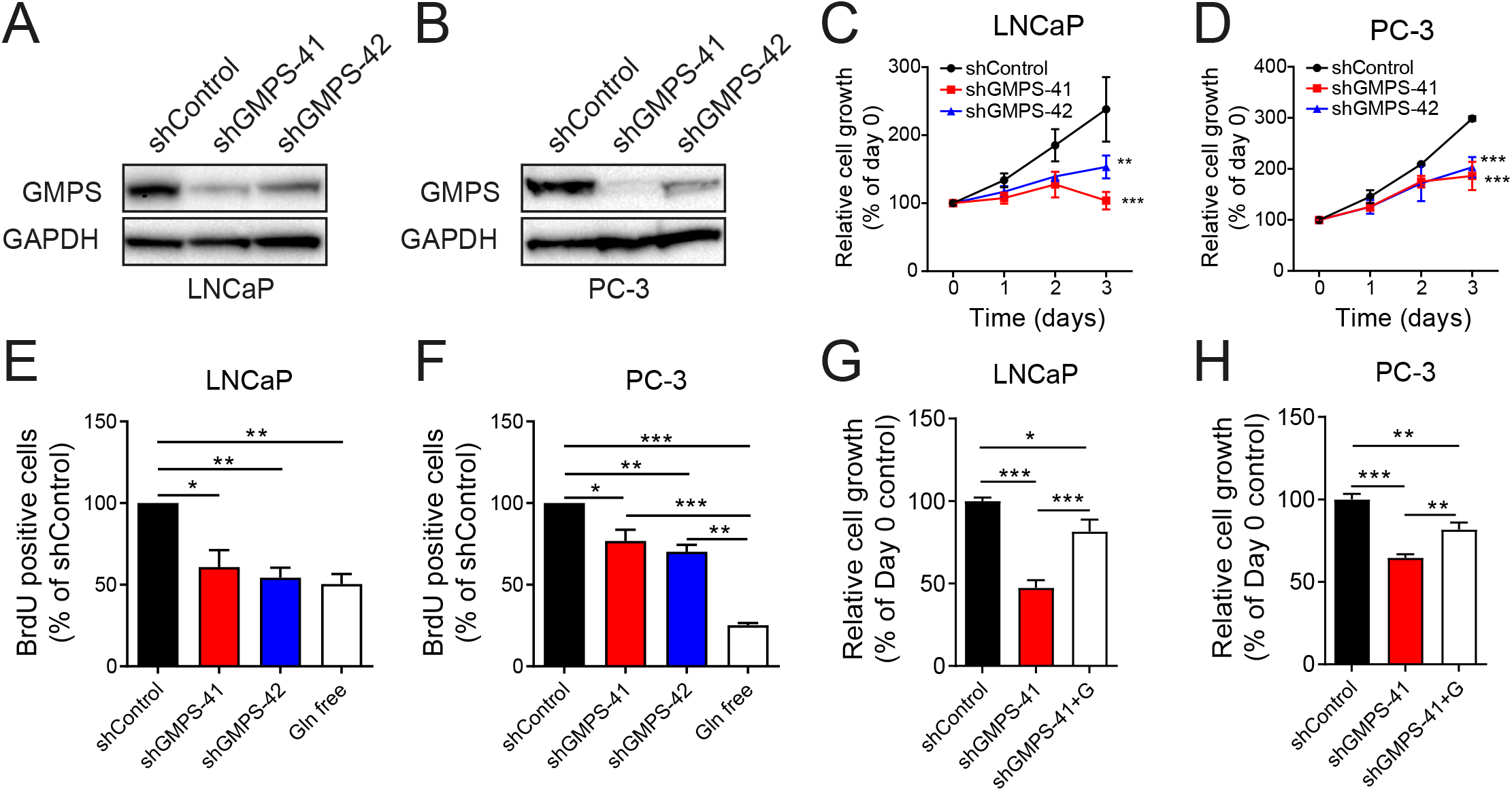
Effect of GMPS knockdown on prostate cancer cells. LNCaP and PC-3 cells were stably transduced with shRNA (shControl or shGMPS-41, shGMPS-42). (A and B) Western blot analysis of GMPS protein after shRNA knockdown in LNCAP (A; n=3) or PC-3 (B; n=3) cells using shGMPS-41 or shGMPS-42. GAPDH was used as loading control. (C and D) Cell growth was assessed in LNCaP (C, n=3) and PC-3 (D, n=3) cells using an MTT assay. Data are mean ± SEM; Two-way ANOVA test. *, P < 0.05; **, P < 0.01; ***, P < 0.001. (E and F) Cell proliferation (BrdU incorporation) was analyzed after shGMPS-41, shGMPS-42 or glutamine (Gln) free media treatment in LNCaP (E; n=3) and PC-3 (F; n=3) cells. (G and H) Exogenous guanosine (200 μM) was added to the culture media, and LNCaP (G) and PC-3 (H) cell growth assessed with or without guanosine over 3 days using an MTT assay. Data are mean ± SEM, n=4. One-way ANOVA test. *, P < 0.05; **, P < 0.01; ***, P < 0.001.

Since GMPS has a number of cellular roles outside of GMP synthesis, we next set out to determine whether activating nucleoside salvage pathways could circumvent GMPS inhibition. LNCaP and PC-3 cells expressing shGMPS were cultured with 200 μM guanosine for three days, which has previously been shown to enable cells to generate GMP through the purine salvage pathway (Bianchi-Smiraglia et al., 2015). In shGMPS cells, cell growth was significantly decreased to 50% and 66% of shControl cells in LNCaP and PC-3 cells, respectively (**Fig. 3G** **and** **3H**). After addition of 200 μM guanosine, cell growth significantly increased to 80% (LNCaP; **Fig. 3G**) and 78% (PC-3; **Fig. 3H**) of shControl cells, suggesting a major mechanism of GMPS inhibition occurs directly through the purine biosynthesis pathway. The inability to completely rescue growth suggested either that the salvage pathways are not sufficient to match normal purine biosynthesis rates, or that other GMPS cellular roles are also important for LNCaP and PC-3 proliferation.

### GMPS knockdown alters purine and pyrimidine biosynthesis from ^15^N-(amide)-glutamine

As guanosine supplementation rescued a significant proportion of cellular proliferation in both LNCaP and PC-3 cells, we next set out to determine the metabolic consequences of GMPS inhibition. Glutamine is a key nitrogen (amide) donor in both purine and pyrimidine nucleotide biosynthesis. In purine biosynthesis, PPAT and PFAS each catalyze the transfer of a glutamine amide within the purinosome to ultimately generate IMP (**Fig. 4A**). These two amide groups are maintained in conversion of IMP to AMP or XMP, with GMPS adding a third glutamine amide to XMP to generate GMP. In pyrimidine biosynthesis, CAD activity adds a glutamine amide group to form carbamoyl phosphate and ultimately UMP, before CTPS1/2 catalyzes addition of a further glutamine amide to form CTP (**Fig. 4A**). In our analysis of the TCGA prostate cancer cohort, we have shown significant positive correlation between GMPS and each of these enzymes (**Fig. S1C**), suggesting they are co-expressed at high levels in higher grade tumors to prioritize purine and pyrimidine biosynthesis. To dissect out how prostate cancer cell lines utilize these pathways, we performed liquid chromatography mass spectrometry (LC-MS) analysis to measure glutamine-derived ^15^N-(amide)-nitrogen contribution to purine and pyrimidine metabolites (**Fig. 4A**). LNCaP and PC-3 cells expressing shControl or shGMPS-41 were cultured in RPMI media containing 2 mM ^15^N-(amide)-glutamine in dialyzed serum (no unlabeled glutamine or free purines and pyrimidines) for 16 h (steady state) and metabolites were extracted for LC-MS analysis. In both prostate cancer cell lines, the majority of glutamine present in the LC-MS analysis was ^15^N labelled as expected (**Fig. 4A**).

**Figure 4.**
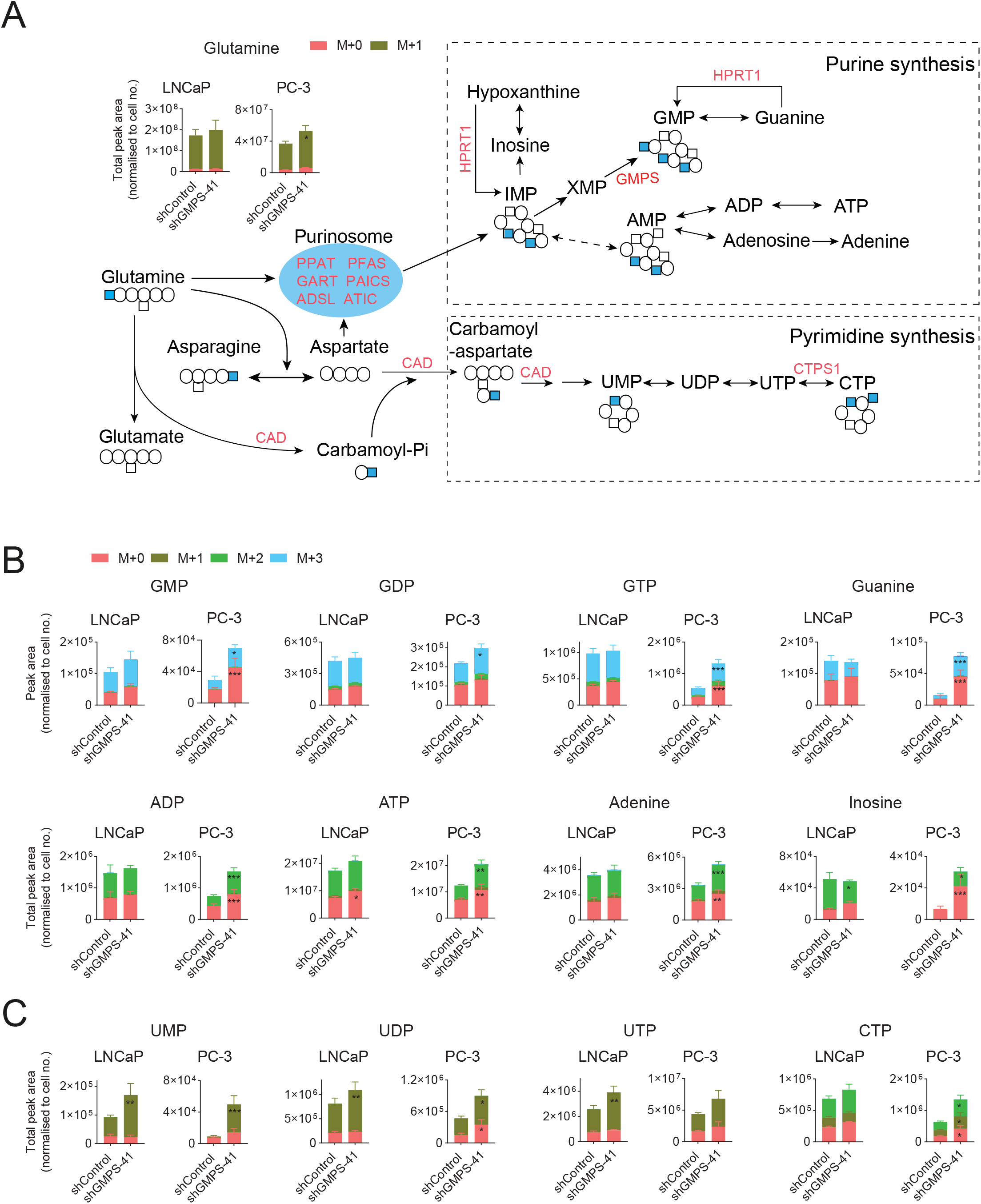
Analysis of ^15^N-(amide)-glutamine metabolites after GMPS knockdown. (A) Schematic showing the contribution of glutamine amide groups to purine and pyrimidine biosynthesis. ^15^N-(amide)-glutamine was added to LNCaP and PC-3 cells stably expressing shRNAs (shControl or shGMPS-41) for 16 h. Abundance of cellular metabolites were measured using LC-MS, with ^15^N-(amide)-glutamine levels showing effective labeling. (B) ^15^N-(amide)-glutamine derived cellular purine metabolites in LNCaP and PC-3 shControl and shGMPS-41 cells. (C) ^15^N-(amide)-glutamine derived cellular pyrimidine metabolites in LNCaP and PC-3 shControl and shGMPS-41 cells. Data are mean ± SEM of 3 independent experiments in duplicates; Two-way ANOVA test. *, P < 0.05; **, P < 0.01; ***, P < 0.001.

A significant increase in the majority of ^15^N-(amide)-glutamine-derived and unlabeled purine metabolites were observed in PC-3 shGMPS cells but not LNCaP cells (**Fig. 4B** **and** **Fig. S3A**). These included GMP, GDP, GTP, guanine, ADP, ATP, adenosine, adenine, inosine, and hypoxanthine (**Fig. 4B** and **Fig. S3A**). These data indicate a global accumulation of purine intermediates in PC-3 shGMPS cells compared to no consistent changes in LNCaP (**Fig. 4B** **and** **Fig. S3A**). While these PC-3 data were surprising, the purinosome has been shown to increase activity at low purine levels (Zhao et al., 2015). To better examine the rate of purine synthesis, we performed additional analyses of ^15^N-(amide)-glutamine metabolomics after a 2 h incubation. As expected, increased ^15^N-amide labeling was seen in shGMPS PC-3 cells in IMP after just 2 h, consistent with increased purinosome activity as an adaptation to low purine levels (**Fig. S3B**). However, despite this increased rate of IMP formation, due to the lower GMPS levels, there was a lower percentage of ^15^N-labeled GMP and guanine in shGMPS cells at 2 h compared to unlabelled GMP and guanine (**Fig. S3C**; GMP:28% shControl, 20% shGMPS; guanine: 21% shControl, 14% shGMPS). The shGMPS cells still express low levels of the GMPS enzyme (**Fig. 3B**), but due to their low proliferation rate (**Fig. 3D**), combined with increased purinosome activity, the cell would reduce the turnover of their purine pool for DNA and RNA synthesis (**Fig. 3F**), thereby leading to higher purine levels.

In addition to increased purinosome activation, we noted higher levels of unlabeled purines in PC-3 cells compared to LNCaP cells (**Fig. 4B**). These unlabeled purines are likely to be generated through the purine salvage pathway. The purine intermediates, guanine and hypoxanthine, can be salvaged and recycled by hypoxanthine-guanine phosphoribosyltransferase (HPRT1) to generate GMP and IMP (**Fig. 4A**). HPRT1 protein is expressed more highly in PC-3 cells compared to LNCaP cells (**Fig. S3D**). Furthermore, HPRT1 expression correlates with GMPS expression in the TCGA cohort (**Fig. S3E**) and HPRT1 is also highly expressed in androgen independent prostate cancer compared to androgen dependent prostate cancer (**Fig. S3F**). This suggests the purine salvage pathway may also be critical in high grade prostate cancer. In PC-3 cells, IMP and GMP contained both unlabeled and labeled pools, with higher amounts of unlabeled GMP (**Fig. 4B**, **Fig. S3C**) and IMP (**Fig. S3B**) in shGMPS cells suggesting increased use of purine salvage.

These two mechanisms of increased synthesis and salvage would allow GMP and other purine intermediates to accumulate in PC-3 shGMPS cells more than LNCaP cells, which would exhibit less salvage pathway, but still maintain less purine usage for cell division. By contrast, purines and pyrimidines in shControl cells will be constantly utilized (and thereby depleted) for DNA and RNA synthesis during cell division. These pathways likely assist in survival of shGMPS cells (low apoptosis; **Fig. S2B, C**), but are not sufficient for driving cell proliferation (**Fig. 3C-F**). The majority of ^15^N-(amide)-glutamine-derived pyrimidine metabolites were significantly increased in shGMPS cells in both PC-3 and LNCaP cell lines (**Fig. 4C** **and** **Fig. S3G**). These included UMP, UDP, UTP and CTP (**Fig. 4C**). Similar to purines, the accumulation of pyrimidines in shGMPS cells likely reflects their lower usage for cell division, although this may also be due to compensatory pyrimidine metabolism or salvage when purine synthesis is disrupted by GMPS knockdown.

### GMPS knockdown alters glutamine-derived carbon usage

Since shGMPS reduces glutamine utilization as an amide donor, we next examined whether this also effects anaplerosis of glutamine carbons using U-^13^C_5_-glutamine, which is ^13^C-labeled at all five glutamine carbons (M+5). Analysis of steady state ^13^C-derived metabolites showed a significant increase in fully labeled (M+5) glutamine, glutamate and α-ketoglutarate (α-KG) retained in shGMPS LNCaP cells. This suggests that while there is an overall reduction in glutamine metabolism after GMPS knockdown, more total glutamine is directed into the TCA cycle through α-KG (**Fig. 5A**). This is confirmed by the increased abundance of downstream TCA metabolites including succinate, fumarate, malate and citrate levels in the LNCaP shGMPS cells (**Fig. 5A**), suggesting that TCA cycle is elevated in GMPS knockdown LNCaP cells. This TCA cycle upregulation appears to be directed toward aspartate accumulation, with a significant increase in aspartate levels in shGMPS cells. Analysis of oxygen consumption rate (OCR) using a Seahorse XF96e bioanalyzer confirmed these metabolomics data, showing a significant increase in the basal and maximal OCR in LNCaP shGMPS cells compared with shControl (**Fig. 5B**). Analysis of extracellular acidification rate (ECAR) showed an increase in LNCaP shGMPS cells (**Fig. 5C**), however there were no significant changes seen in either pyruvate or lactate levels after shGMPS knockdown (**Fig. 5A**).

**Figure 5.**
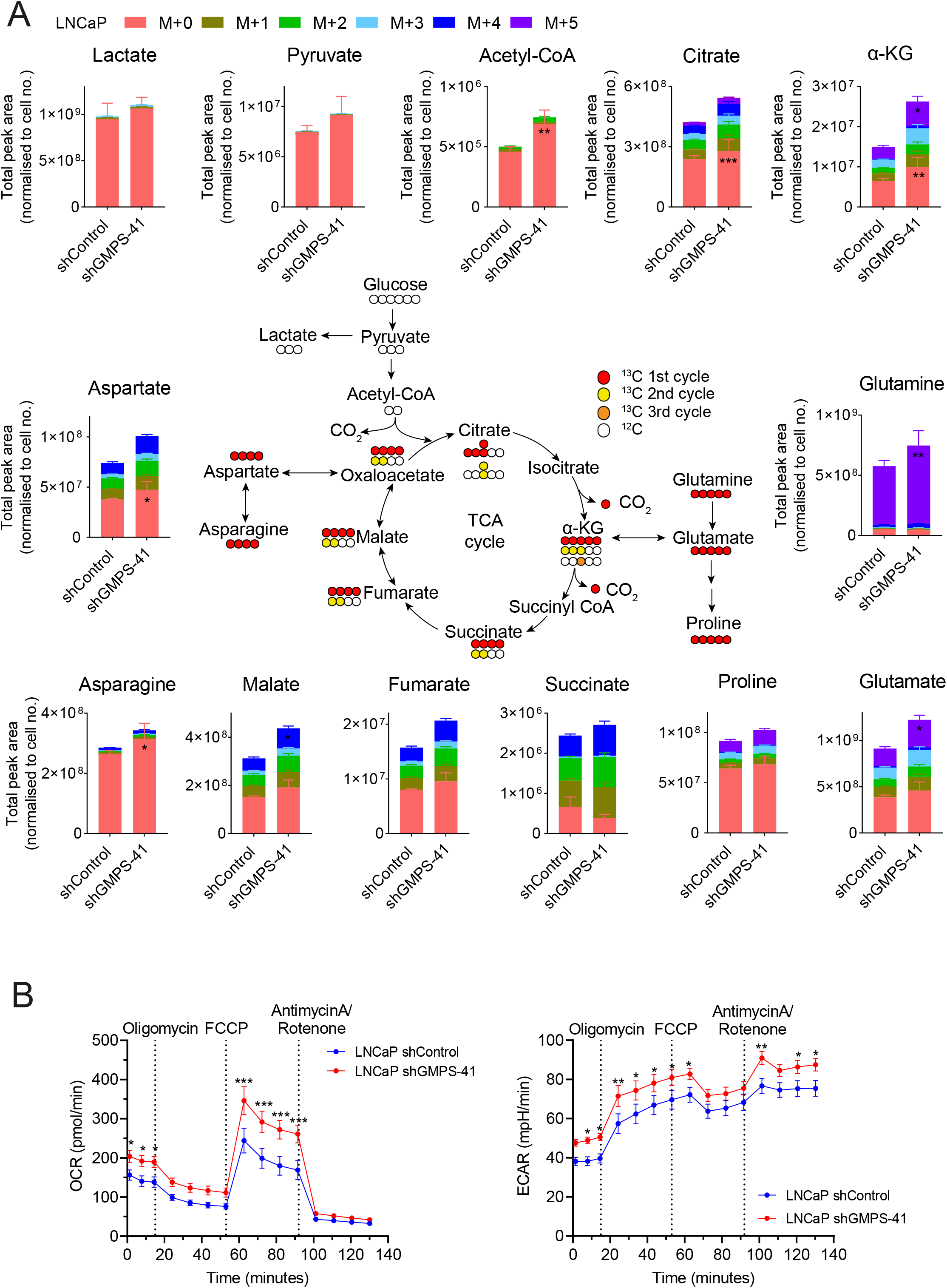
Glutamine-derived carbon usage in LNCaP cells after GMPS knockdown. (A) Schematic showing the contribution of glutamine carbon to TCA Cycle. U-^13^C_5_-glutamine was added to LNCaP stably expressing shRNAs (shControl or shGMPS-41) for 16 h. Abundance of cellular metabolites were measured using LC-MS. Mass isotopologue distribution of U-^13^C_5_-glutamine derived TCA metabolites in LNCaP shControl and shGMPS-41 cells are shown. Data are mean ± SEM of 3 independent experiments in duplicates; Two-way ANOVA test. *, P < 0.05; **, P < 0.01; ***, P < 0.001. (B and C) Oxygen consumption rate (OCR; B) and extracellular acidification rate (ECAR; C) were measured using Mito Stress Test kit on a Seahorse XF96e analyzer in LNCaP shControl and shGMPS-41 cells. Data are mean ± SEM, n=5; Two-way ANOVA test. *, P < 0.05; **, P < 0.01; ***, P < 0.001.

By contrast, shGMPS in PC-3 cells showed no consistent changes in glutamine, glutamate or α-KG levels (**Fig. 6A**). Malate and fumarate levels (M+4) were significantly decreased in PC-3 shGMPS cells, while there was a trend to decreased succinate with no change in citrate levels (**Fig. 6A**). Interestingly, there was a significant increase in aspartate levels from both glutamine (M+4) and non-glutamine sources (M+0) in PC-3 shGMPS cells (**Fig. 6A**). Aspartate is involved in malate-aspartate shuttle and can be synthesized from TCA intermediate oxaloacetate (OAA) by GOT2. The increased glutamine-derived aspartate levels suggest that more M+4 metabolites leave TCA cycle and are stored as aspartate in PC-3 cells, thereby depleting the TCA cycle metabolite pool. Despite the divergent TCA cycle mechanisms for LNCaP and PC-3 cells, both accumulate significantly higher aspartate levels. While this may be directly related to changes the TCA cycle, it is also possible the high aspartate levels are due to decreased aspartate catabolism through the purine and pyrimidine pathways (**Fig. 4A**).

**Figure 6.**
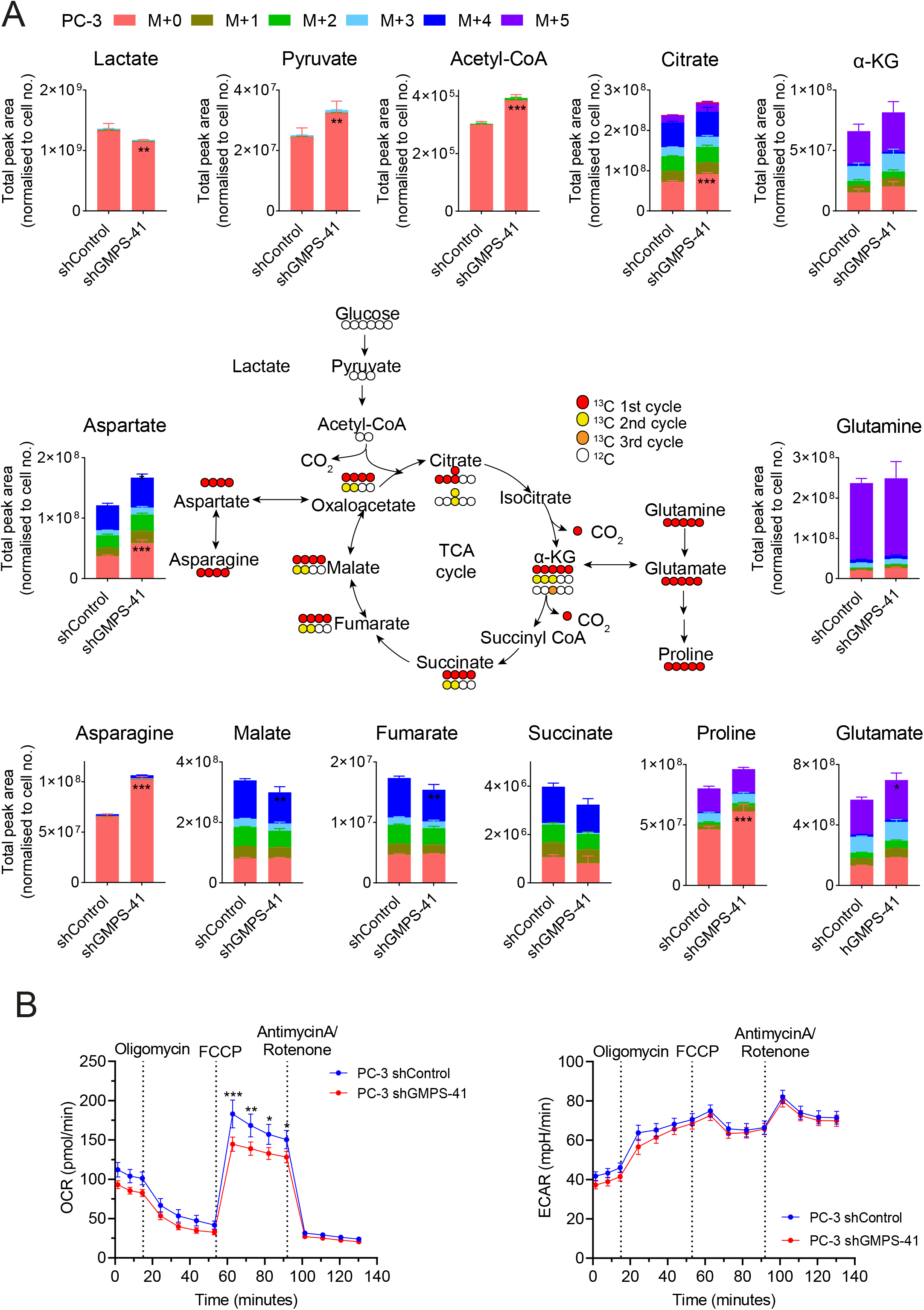
Glutamine-derived carbon usage in PC-3 cells after GMPS knockdown. (A) Schematic showing the contribution of glutamine carbon to TCA Cycle. U-^13^C_5_-glutamine was added to PC-3 cells stably expressing shRNAs (shControl or shGMPS-41) for 16 h. Abundance of cellular metabolites were measured using LC-MS. Mass isotopologue distribution of U-^13^C_5_-glutamine derived TCA metabolites in PC-3 shControl and shGMPS cells are shown. Data are mean ± SEM of 3 independent experiments in duplicates; Two-way ANOVA test. *, P < 0.05; **, P < 0.01; ***, P < 0.001. (B and C) Oxygen consumption rate (OCR; B) and extracellular acidification rate (ECAR; C) were measured using Mito Stress Test kit on a Seahorse XF96e analyzer in PC-3 shControl and shGMPS-41 cells. Data are mean ± SEM, n=5; Two-way ANOVA test. *, P < 0.05; **, P < 0.01; ***, P < 0.001.

We further measured the metabolites of glycolysis, including lactate, pyruvate and acetyl-CoA, which are mainly unlabeled metabolites likely derived from glucose (**Fig. 6A**). Acetyl-CoA is a versatile metabolite that serve as a precursor in many metabolic pathways, a significant increase in acetyl-CoA levels were found in both LNCaP and PC-3 shGMPS cells (**Fig. 5A** **and** **6A**). In PC-3 cells, pyruvate levels were also significantly increased, while lactate levels were significantly decreased (**Fig. 6A**). These results suggested that aerobic glycolysis is suppressed, and more glucose-derived carbon enters TCA cycle when GMPS is knocked down in PC-3 cells, with more of these carbons also likely to exit as aspartate to support purine and pyrimidine metabolism, as shown by the significant increase in unlabeled aspartate (M+0) (**Fig. 6A**). PC-3 cells overall showed lower oxygen consumption rate, with a significant decrease in maximal OCR in shGMPS cells (**Fig. 6B**), however there was no change in the ECAR despite significantly lower lactate levels (**Fig. 6C**). Together these data suggest that when GMPS is knocked down, cells are able to reprogram their metabolism and utilize alternative pathways to generate energy and accumulate required metabolite pools for survival, albeit at a significantly reduced proliferative capacity.

### GMPS knockdown inhibits PC-3 xenograft tumor growth

To determine whether GMPS function is necessary for tumor growth *in vivo*, PC-3 cells expressing shControl or shGMPS-41 were subcutaneously injected into nude mice. Knockdown of GMPS had no effect on the tumor take rate: 65% for shControl compared with 60% for shGMPS cells. Tumor volume was measured twice per week for 28 days, showing a significant decrease in shGMPS tumor size by day 20 (**Fig. 7A**). Mice were euthanized after 28 days at ethical endpoint due to the size of the shControl tumors. The tumors were isolated, photographed and weighed, with shGMPS tumors showing a significant reduction in weight and were visually smaller compared to shControl tumors (**Fig. 7B** **and** **7C**). Immunofluorescence staining analysis of the tumors showed that expression of the proliferation marker Ki67 was consistently lower in the shGMPS tumors compared to shControl tumors (**Fig. 7D**). These data suggested that GMPS plays an important role in prostate cancer growth *in vivo* via inhibition of proliferation.

**Figure 7.**
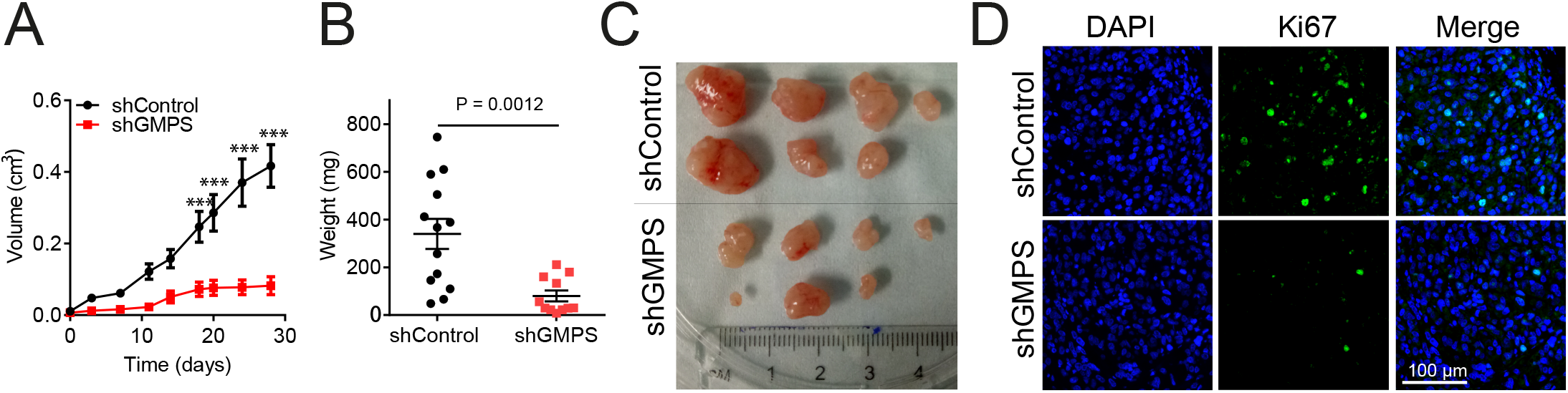
GMPS knockdown blocks tumor progression in PC-3 xenografts in vivo. (A) PC-3 cells stably expressing shRNAs (shControl or shGMPS-41) were implanted subcutaneously into the left and right flank of 8-12-week-old male athymic nu/nu mice. Tumor growth was monitored using calipers. Data are mean ± SEM, n=13 for shControl; n=11 for shGMPS; Two-way ANOVA test. (B) Weight of tumors was measured. Data are mean ± SEM, n=13 for shControl; n=11 for shGMPS; Two-way ANOVA test. (C) Photo of representative tumors for shControl and shGMPS after harvesting. (D) Cell proliferation marker Ki67 was detected by immunofluorescence in shControl and shGMPS PC-3 xenograft tumors. Scale bar is 100 μm. *, P < 0.05; **, P < 0.01; ***, P < 0.001.

## Discussion

In this study, we demonstrated the importance of increased GMPS expression in advanced prostate cancer and its function in the supply of glutamine amide groups for purine biosynthesis, further defining the metabolic outcomes of glutamine addiction in prostate cancer. While it has been well defined that glutamine uptake mediated by ASCT2 is critical in prostate cancer cells (Wang et al., 2015), our study provides a new therapeutic target (GMPS) and understanding of how glutamine is metabolized and utilized in both androgen dependent and independent cell lines. We show not only that GMPS is significantly upregulated in prostate cancer samples, but that its expression significantly correlates with the major enzymes involved in producing purines and pyrimidines from glutamine. In addition, we show that GMPS protein is highly expressed in a cohort of patient samples, as is the glutamine transporter ASCT2, further backing up these mRNA data. These data suggest that at least a proportion of prostate cancer cells are wired with all the downstream machinery to efficiently prioritize purine and pyrimidine biosynthesis, as well as the mechanism to uptake glutamine into the cells. Furthermore, these high GMPS expressing prostate cancers were more frequently of higher grade and were more metastatic, resulting in significant decreases in both disease/progression-free and overall survival.

Recent data has shown the importance of glutamine metabolism in neuroendocrine prostate cancer (Mishra et al., 2018). Mishra *et al* showed that glutamine could be provided to the cancer cells by cancer-associated fibroblasts, which was critical for metabolic reprogramming of the cancer cells to become more dependent on glutamine and drive neuroendocrine differentiation (Mishra et al., 2018). In that study, blocking ASCT2-mediated glutamine uptake reduced tumor size in a CRPC model, but only glutamine carbons were traced into TCA cycle, and it is unclear whether glutamine was also utilized for nucleotide synthesis in these models (Mishra et al., 2018). Glutamine metabolism has also been shown as a vulnerability in a metastatic subline of PC-3, whereby glutaminase inhibition becomes more effective (Zacharias et al., 2017). This PC-3M subline also showed increased glutamine carbon flux into the TCA cycle (Zacharias et al., 2017). In addition, there are differences between glutamine carbon use in different prostate cancer cell lines, with LNCaP cells previously shown to prioritize lipid synthesis from glutamine carbons when ASCT2-mediated glutamine uptake was blocked (Wang et al., 2015). A common thread throughout these studies is that prostate cancer cells are generally reliant on glutamine, a dependency which becomes more apparent in aggressive tumors. Our glutamine carbon tracing data confirm previous reports on the important contribution of glutamine to TCA cycle, with ~56% of the carbons being derived from glutamine within TCA cycle metabolites in LNCaP cells, ~74% of the carbons being derived from glutamine in PC-3 shControl cells. Interestingly, in LNCaP cells, the labeled proportion increased after GMPS knockdown, with a ~28% increase in the ^13^C-labeled:unlabeled ratio in TCA metabolites in LNCaP cells (~5% decrease in PC-3 cells). In addition, our ^15^N-amide tracing data clearly shows for the first time the critical role for glutamine in producing both purines and pyrimidines in prostate cancer cell lines. Over a 16 h pulse, glutamine amide-labeling accounted for ~44% (PC-3) and ~59% (LNCaP) of the total pool of purine metabolites analyzed and ~63% (PC-3) and ~71% (LNCaP) of the total pool of pyrimidine metabolites analyzed. These data also show the increased use of the purine salvage pathway in PC-3 cells compared to LNCaP.

The *de novo* biosynthesis pathway of GMP is an energy intensive process, requiring three glutamines, six ATPs, two formates, one glycine and one aspartate to generate a single GMP (Pedley and Benkovic, 2017). The salvage pathway is more economical to maintain the purine pool because HPRT1 can recycle guanine and hypoxanthine to generate GMP and IMP with a much reduced energy requirement (Yin et al., 2018). IMP can be synthesized to AMP or GMP again. Our data showed that PC-3 cells are more dependent on salvage pathway comparing to LNCaP cells, therefore less energy is required and oxygen consumption rate is decreased in PC-3 cells. Knockdown of GMPS delayed the *de novo* synthesis of GMP in prostate cancer cells and suppressed cell proliferation, leading to accumulation of purine and pyrimidine in cells. Although the salvage pathway provides more IMP and GMP for purine synthesis, the total purine pool was still lower than shControl cells due to their high rate of cell division. Despite an active nucleotide salvage pathway in PC-3 cells facilitating cell survival, GMPS remains essential for rapid cell proliferation.

GMPS expression is high in other cancers such as melanoma and ovarian cancer (Bianchi-Smiraglia et al., 2015; Wang et al., 2019). GMPS expression is increased in metastatic melanoma and was recently shown to be a prognostic marker of ovarian cancer progression (Bianchi-Smiraglia et al., 2015; Wang et al., 2019). Here we show that GMPS inhibition by either pharmacologic or shRNA knockdown blocks tumor growth *in vitro* and *in vivo*, similar to what was shown for melanoma cells (Bianchi-Smiraglia et al., 2015). In addition to these roles in facilitating proliferation, there appears to be a putative role for GMPS in invasion, seen through both melanoma functional studies (Bianchi-Smiraglia et al., 2015) as well as colocalization studies in renal cancer, which suggest a purine biosynthesis compartment is present at the leading edge of cancer cells (Wolfe et al., 2019).

Overall, our metabolomics data have revealed substantial changes in pyrimidine metabolite levels in both LNCaP and PC-3 in response to GMPS knockdown. Since guanosine addition was able to rescue the growth phenotype, this pyrimidine accumulation is likely due to the inhibition of cell cycle resulting in additional pyrimidines that are not required for DNA replication. The increased purine metabolite levels in PC-3 cells, but not LNCaP cells, are intriguing. The guanosine rescue experiment resulted in a more complete rescue of cell growth in LNCaP cells, which may result from differences in the efficiency of the purine salvage pathway between these two cell lines, consistent with the higher expression of HPRT1 protein in PC-3 cells. The TCA cycle alterations present in LNCaP cells and PC-3 cells, while causing distinct metabolite patterns, both resulted in increased aspartate, a critical purine and pyrimidine precursor. These changes in TCA cycle were evident both in glutamine and non-glutamine carbons, and may also be fueled by fumarate production, which is also a product of multiple steps in purine biosynthesis. This may provide an alternative explanation as to why increases were observed in fumarate and malate in LNCaP cells, but not in upstream succinate, ultimately resulting instead in increased levels of aspartate in a futile attempt to replenish purine biosynthesis.

In addition to the metabolic role of GMPS, it has been reported that genotoxic stress or nucleotide deprivation triggers accumulation of GMPS in the nucleus (Reddy et al., 2014). This nuclear accumulation facilitates GMPS interaction with USP7 (van der Knaap et al., 2010; van der Knaap et al., 2005), a protein which plays a role in stabilizing p53 and thereby activating its transcription (Reddy et al., 2014). USP7/HAUSP is also increased in prostate cancer and has been shown to regulate PTEN nuclear exclusion in aggressive prostate cancer (Song et al., 2008). Our data show substantial GMPS expression in the nucleus and cytoplasm both in cell lines and patient samples, suggesting this nuclear role may be important in addition to its metabolic role, particularly since prostate cancers frequently contain p53 alterations (Cancer Genome Atlas Research, 2015; Carroll et al., 1993; Dahiya et al., 1996; Ecke et al., 2010). Gain of function p53 mutants are important for androgen-independent growth of prostate cancer cells (Nesslinger et al., 2003; Vinall et al., 2006), and it is also possible that GMPS may affect other targets of USP7, such as histone H2B, which requires further study.

Overall, our study has shown that high GMPS expression and enzymatic function is critical for prostate cancer cell proliferation and tumor growth. Furthermore, this function is an important component of glutamine addiction in prostate cancer cells, suggesting that GMPS is a putative therapeutic target with a potential companion biomarker.

## Materials and Methods

### Patient Samples

A cohort of 64 patients with prostate cancer was used in this study. Patients underwent a radical prostatectomy (RP) for localized prostate cancer by a specialist urologist at St. Vincent’s Hospital, Sydney. This project was approved by the St. Vincent’s Campus Research Ethics Committee (Approval No. 12/231) and informed consent was obtained from all human participants. Tissue microarrays (TMA) containing 1.5 mm cores of prostate adenocarcinoma were constructed from formalin-fixed paraffin-embedded RP tissue blocks as previously described (Horvath et al., 2005). Each patient case was represented by a mean of 3 biopsies (range, 2–5 biopsies) of prostate cancer of different primary Gleason score and one biopsy of hyperplasia adjacent to cancer, with a total of 63 cases containing enough tissue for the immunohistochemistry analyses.

### Immunohistochemistry

Immunostaining of TMA sections (4 μM thickness) with a GMPS monoclonal mouse antibody (1:250, Santa Cruz, sc-376163) or ASCT2 rabbit antibody (1:2000, Sigma, HPA035240) was performed on a Leica Bond automated stainer, with heat induced epitope retrieval ER2 for 30 min, primary incubation for 60 min, and detection system with the Bond Refine Polymer kit. Immunostaining was scored by staining intensity (graded as 0 [absent], 1 [weak], 2 [moderate], or 3 [strong]) along with the percent of cells stained.

### Cell lines

Human prostate cancer cell lines LNCaP-FGC, PC-3 and DU145 were purchased from ATCC (Manassas, VA, USA). We confirmed LNCaP, PC-3, DU145 cell identity by STR profiling (Cellbank, Australia). Cells were cultured in RPMI 1640 medium (Invitrogen, Australia) containing 10% (v/v) fetal bovine serum (FBS), penicillin-streptomycin solution (Sigma-Aldrich, Australia) and 1 mM sodium pyruvate (Invitrogen, Australia). Cells were maintained at 37°C in a fully humidified atmosphere containing 5% CO_2_.

### Lentiviral shRNA expression

GMPS shRNA lentiviral preparation was performed as previously described (Wang et al., 2014). Two different shRNAs for GMPS were used in this study (Sigma), shGMPS-41: CCGGGCATTTGCTATAAAGGAACAACTCGAGTTGTTCCTTTATAGCAAATGCTTTTTG; shGMPS-42: CCGGCCTACAGTTACGTGTGTGGAACTCGAGTTCCACACACGTAACTGTAGGTTTTTG. Briefly, pLKO.1 plasmid containing short hairpin RNA (shRNA) against GMPS or a non-targeting *Arabidopsis thaliana* miR159a control sequence (shControl) were mixed with pMDLg/prre, pRSVRev and pMD2.VSV-G packaging plasmids, and transfected into 70% confluent HEK293T cells using the calcium phosphate precipitation method. After 8 h, the media was changed to fresh RPMI media containing 25 μM chloroquine. Viral supernatant was collected 24 h later and concentrated by ultracentrifugation, then snap-frozen for storage at −80°C until use. Viral aliquots were used to transduce LNCaP and PC-3 cells in the presence of 8 μg/ml polybrene. These cells were expanded under puromycin selection (10 μg/mL) for at least 1 week. GMPS expression was examined by western blotting to confirm knockdown.

### Western blots

Cells were seeded at a density of 0.5 × 10^6^ in 10 cm plates, allowed to adhere overnight. Cells were lyzed by addition of lysis buffer with Protease Inhibitor Cocktail III (Bioprocessing Biochemical, California) and Phosphatase Inhibitor Cocktail (Cell Signaling Technology). Equal amount of protein (micro-BCA method; Pierce, IL) were loaded on 4-12% gradient gels (Invitrogen, California), electrophoresed and transferred to PVDF membrane. The membrane was blocked with 2.5% (w/v) BSA in PBS-Tween 20, and incubated with the primary antibodies, washed and incubated with secondary antibodies. After washing, the secondary HRP-labelled antibodies were detected using enhanced chemiluminescence reagents (Pierce) on a ChemiDoc Imager (BioRad). Antibodies used were mouse IgG against GMPS (Santa Cruz), rabbit IgG against HPRT1 (Abcam), rabbit IgG against cleaved-PARP (Cell Signaling Technologies) or a mouse IgG against glyceraldehyde-3-phosphate dehydrogenase (GAPDH; Abcam). Horseradish peroxidase-conjugated donkey anti-mouse IgG, and donkey anti-rabbit IgG were used as secondary antibodies (Millipore).

### Cell viability assay

Cells in exponential growth phase were harvested and seeded (1 × 10^4^/well) in a flat-bottomed 96–well plate. The cells were incubated overnight in RPMI media, prior to culture with or without 200 μM Decoyinine (Deco; AdipoGen Life Sciences) or glutamine free media for 24, 48 or 72 h. MTT solution (10 μL; 3–(4,5–dimethylthiazol–2–yl)–2,5–diphenyl tetrasodium bromide; Millipore) was added to each well for 4 h, prior to addition of 100 μL of isopropanol/HCl solution and mixed thoroughly. Plates were read at 570 nm and 630 nm in a plate reader Infinite 200 Pro (TECAN, Switzerland). Results were plotted as percentages of the absorbance observed in control wells.

### BrdU analysis

Cells (2 × 10^5^ per well) were seeded in 6-well plates and allowed to adhere overnight in serum-free media. After serum starvation, cells were incubated in the fresh RPMI media for 22 h, followed by addition of BrdU (150 μg/mL) for another 2 h. Cells were detached using Tryple (Life Technologies), fixed and stained using the APC-BrdU Flow Kits (BD, Biosciences). The BrdU antibody was diluted 1:50. Nuclei were counter-stained with 7-AAD. BrdU incorporation analysis was performed using a BD Canto II flow cytometer and data was analyzed by FlowJo software (Tree Star Inc.).

### Annexin V assay

Cells (2 × 10^5^ per well) were seeded in 6-well plates, allowed to adhere overnight. Cells were treated with decoyinine, or cultured in normal media or glutamine free media for 48 h. Cells were detached using TrypLE and resuspended in 1 mL of binding buffer (HEPES-buffered PBS supplemented with 2.5 mM calcium chloride) containing anti-annexin V-APC (BD) and incubated for 15 min in the dark at room temperature. Propidium iodide (PI) solution (20 μg/mL) was added, and the cells were analyzed using a BD Canto II flow cytometer and FlowJo software (Tree Star Inc.).

### Immunofluorescence staining

Cells were seeded on coverslips at a density of 0.5 × 10 ^6^ cells/well in a 6-well plate and allowed to adhere overnight. After serum starvation for 2 h, the cells were incubated with fresh media for 24 h. Cells were fixed using 4% (w/v) paraformaldehyde (Thermo Fisher Scientific) for 20 min, and permeabilization using 0.1% PBST for 15 min. Cells were washed and incubated with 5% (v/v) normal goat serum in 2% (w/v) BSA/PBS for 30 min before addition of the GMPS monoclonal antibody (1:200 dilution) at 4°C for overnight incubation. The cells were washed in PBS before addition of a goat anti-mouse or anti-rabbit IgG conjugated with AlexaFluor 488 (1:1000; Thermo Fisher Scientific) for 1 h at room temperature, and nuclear staining with DAPI (1:1000) for 5 min. Cells were washed in PBS, coverslips immersed in glycerol, placed on a slide and visualized using a Leica DM6000B (Leica).

### Seahorse bioanalyzer

Cellular oxygen consumption rate (OCR) and extracellular acidification rate (ECAR) was measured with a Seahorse XF96e bioanalyzer (Seahorse Bioscience, MA) using a Cell Mito Stress Test Kit according to manufacturer’s instructions. Briefly, cells (2 × 10^4^ cells/well) were seeded in a Seahorse XF 96-well assay plate in full growth medium. After overnight attachment, the medium was carefully washed and replaced with pre-warmed running medium (consisting of non-buffered DMEM (Agilent Technologies, CA) supplemented with 1 mM sodium pyruvate, 2 mM glutamine and 10 mM glucose, pH 7.4). Plates were incubated for 60 min in a non-CO_2_ incubator at 37°C before three basal measurements were undertaken determining oxygen and proton concentration in the medium. The ATP synthase inhibitor oligomycin (1 μg/mL) was immediately added followed by four further measurements; Carbonyl cyanide-*4*-(trifluoromethoxy) phenylhydrazone (FCCP; 0.5 μM) was then added with another four further measurements, before addition of the complex I and III inhibitors rotenone/antimycin A (0.5 μM) and a final four further measurements taken.

### *In vitro* L-^15^N-(amide) glutamine and L-U-^13^C_5_ glutamine tracing and metabolite extraction

For 2 h and steady-state (16 h) labeling, cells were seeded in 6-well plates in duplicate from three different passages at a density of 0.5 × 10^6^ cells/well in RPMI-1640, with an extra well seeded for cell counts at the time of metabolite extraction for normalization. After 24 h, the cells were washed once with PBS prior to media change to glutamine-free RPMI containing 10% dialyzed fetal bovine serum (Thermo Fisher Scientific), and 2 mM L-^15^N-(amide)-glutamine or L-U-^13^C_5_-glutamine (Sigma). After 2 h or 16 h, metabolites were extracted from cells. After removal of growth media cells were washed three times with 1 mL ice-cold PBS and rapidly lyzed with 1 mL ice-cold extraction buffer (methanol: acetonitrile: MilliQ water at 50:30:20 v/v) at 1 mL/1 × 10^6^ cells. Cells were scraped in the extraction buffer and then transferred into pre-chilled Eppendorf tubes. Blank wells with no cells were also processed as above for background subtraction. The mixtures were then incubated at 4°C for 10 min on a rotator, followed by centrifugation at 14,000 RPM at 4°C for 10 min. The supernatant was collected and used for LC - MS analysis.

### Targeted metabolomics by LC-MS

Metabolomics analysis was performed as described by Mackay et al (Mackay et al., 2015). Briefly, a Q Exactive HF mass spectrometer (Thermo Fisher Scientific) was used with a resolution of 120,000 at 200 mass/charge ratio *(m/z)*, mass scan range of 75 to 1000 *m/z* were used, coupled with a Thermo Ultimate 3000 high-performance liquid chromatography (HPLC) system. For the LC-MS method, a 150 × 2.1 mm, 5 μm SeQuant ZIC-pHILIC column (Merck, VIC, Australia), with a 20 × 2.1 mm SeQuant ZIC-pHILIC guard column was used. Metabolite extracts were injected (4 μL), and compounds were separated using an aqueous mobile phase gradient started with 20% of 20 mM ammonium carbonate adjusted to pH 9.4 with 0.1% ammonium hydroxide and an organic mobile phase of 80% acetonitrile. A linear gradient from 80% organic to 80% aqueous gradient ran for 17 min, followed by an equilibration back to the starting conditions. The flow rate was 200 μL/min with the column oven temperature maintained at 45°C, for a total run time of 26 min. All metabolites were detected using lock masses and mass accuracy was below 5 ppm. Metabolite identification was predetermined with in-house standards where available and isotope labelling patterns of each metabolite were analyzed by Xcalibur (v4.1, Thermo Fisher Scientific). Mass of the isotopologues were corrected by deducting the naturally occurring stable carbon and nitrogen isotopes.

### PC-3 xenografts

Athymic (nu/nu) male nude mice (Animal Resource Center, Perth, Australia) 6-8 weeks of age were housed in a specific pathogen-free facility in accordance with the University of Sydney animal ethics committee guidelines (Approval 2016-021). Mice were anaesthetized via 2% isoflurane inhalation and received subcutaneous (s.c.) injections of 2 × 10^6^ PC-3 cells with shControl or shGMPS-41 resuspended in 100 μL of Hank’s Balanced Salt solution (HBSS). Xenografts were transplanted in both the right and left side dorsal flanks of mice as detailed previously (Wang et al., 2015), using five mice per group from two independent experiments. Tumor growth was monitored via caliper measuring and performed twice a week thereafter for 28 days. Tumor volume was calculated using the formula Volume = Length × Width ^2^×π/6. After 28 days, animals were sacrificed following the final measuring point. After being imaged and weighed, tumors were collected and fixed in 10% (v/v) neutral buffered formalin for embedding and sectioning. For PC-3 xenografts, sections were boiled in sodium citrate buffer (10 mM, pH=6) for 20 min, followed by a rinse in distilled water. After incubation with 5% (v/v) normal goat serum for 30 min, sections were stained for Ki67 antibody (1:100 dilution; Abcam) overnight at 4°C. The cells were washed in PBS before addition of a goat anti-rabbit Alexa 488 (1:1000; Thermo Fisher Scientific) for 2 h at room temperature. After washing with PBS, cells were counterstained with DAPI (Thermo Fisher Scientific) and slides were mounted by glycerol. Staining results were imaged using a Leica DM6000B (Leica).

### Bioinformatics and Statistical analysis

Expression data were generated using online datasets as indicated. TCGA expression and survival data were analyzed using cBioportal (TCGA, Firehose Legacy dataset). All experimental data are expressed as mean ± SEM and were performed using at least 3 replicate experiments. Statistical analysis was performed using a Mann-Whitney U-test or one-way ANOVA test, MTT assay, tumor growth curve analysis and isotopologues analyses used a two-way ANOVA test. All statistical tests were performed in in GraphPad Prism v8 using two-sided tests unless otherwise stated, with p<0.05 considered significant.

## Acknowledgements

This work was supported by grants from Movember through the Prostate Cancer Foundation of Australia (YI0813 to Q. Wang), Tour de Cure (J. Holst) and the Australian Movember Revolutionary Team Award Targeting Advanced Prostate Cancer (MRTA1), J. Holst, and Q. Wang).

## Authors’ Contributions

Conception and design: Q. Wang, Y. F. Guan, J. Holst

Development of methodology: Q. Wang, Y. F. Guan, B. Mak, S. E. Hancock, A. Pang, J. Holst

Acquisition of data: Q. Wang, Y. F. Guan, R. Nagarajah, B. Mak, S. E. Hancock, A. Pang, K. Wahi, M. van Geldermalsen, B. K. Zhang,

Analysis and interpretation of data: Q. Wang, Y. F. Guan, B. Mak, S. E. Hancock, A. Pang, N. Turner, L. G. Horvath, J. Holst

Writing, review, and/or revision of the manuscript: Q. Wang, Y. F. Guan, R. N. Turner, L. G. Horvath, J. Holst

---

**Fig. S1:**
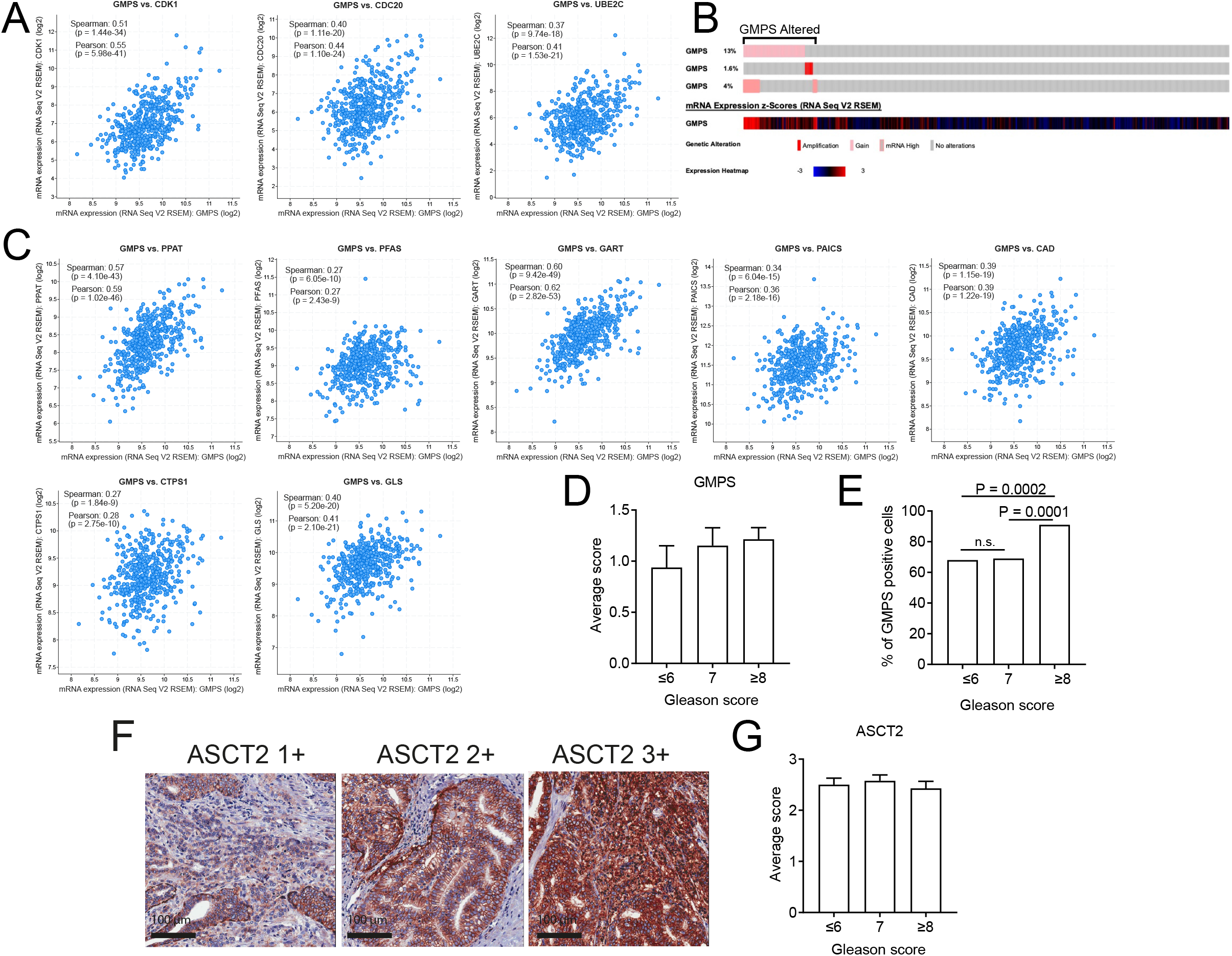
A, GMPS mRNA exprssion correlates with metastatic castration resistant prostate cancer cell cycle genes CDK1, CDC20 and UBE2C in the TCGA dataset (cBioPortal). B, GMPS expression was assessed in TCGA prostate cancer Firehose dataset for GMPS gain, amplification and mRNA expression z-score ≥2. Altered samples were used to generate Kaplan Meier curves in Figure 1. C, GMPS expression correlates with purine (PPAT, PFAS, GART and PAICS) and pyrimidine (CAD and CTPS1) synthesis and glutaminolysis enzymes (GLS) in the TCGA dataset. D,Scoring of GMPS expression in our patient cohort with different Gleason grades (≤6, n = 16; 7, n = 33; ≥8, n = 14). Data are mean ± SEM. One-way ANOVA test. E, Quantification of GMPS protein expression in prostate cancer patient samples from Gleason grade ≤6 (n = 16), Gleason grade 7 (n = 33), Gleason grade ≥8 (n = 14). Data are mean ± SEM. Two tailed Fisher’s exact test. (F) Representative images of ASCT2 protein expression by immunohistochemistry in prostate cancer patient samples. Scale bar is 100 μm. (G) Scoring of ASCT2 expression in patient cohort with different Gleason grades (≤6, n = 16; 7, n = 33; ≥8, n = 14). Data are mean ± SEM.

**Fig. S2:**
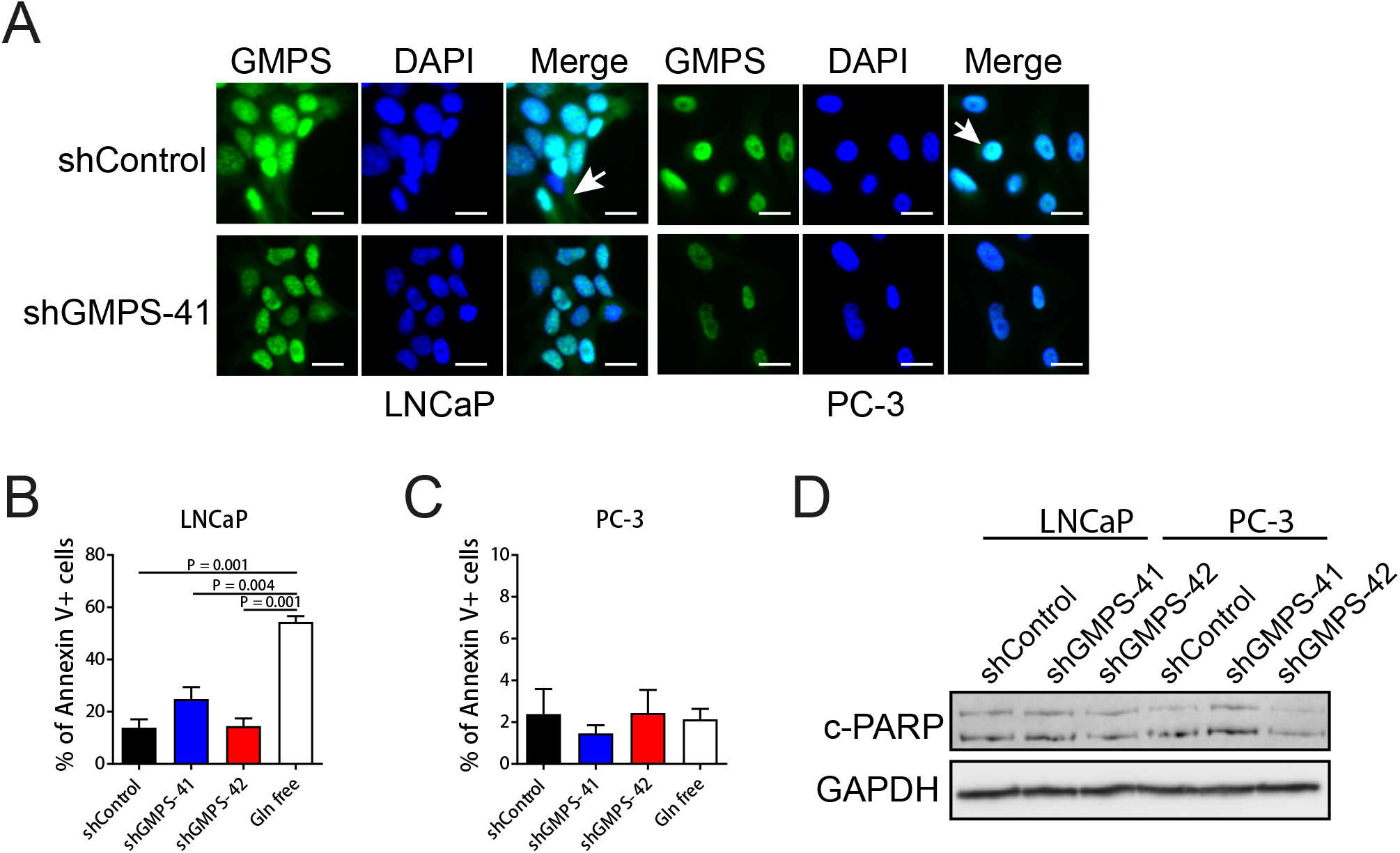
A, Localization of GMPS protein in LNCaP and PC-3 cells was determined by immunofluorescent staining, with nuclei visualized using 40,6-diamidino-2-phenylindole (DAPI). Scale bar is 20μm. B, LNCaP shControl, shGMPS-41 and shGMPS-42 cells were cultured in RPMI media or LNCaP shControl cells in Gln free media for 48 h. Cells were stained using an Annexin-V antibody and PI, and examined by flow cytometry.Data are mean±SEM. n=3. One-way ANOVA test is performed. B, PC-3 shControl, shGMPS-41 and shGMPS-42 cells were cultured in RPMI media or PC-3 shControl cells in Gln free media for 48 h. Cells were stained using an Annexin-V antibody and PI, and examined by flow cytometry.Data are mean±SEM. n=3. One-way ANOVA test is performed. D, Western blot analysis of c-PARP after GMPS knockdown using shGMPS-41or shGMPS-42 in LNCAP and PC-3 cells. GAPDH is used as loading control. n=3.

**Fig. S3:**
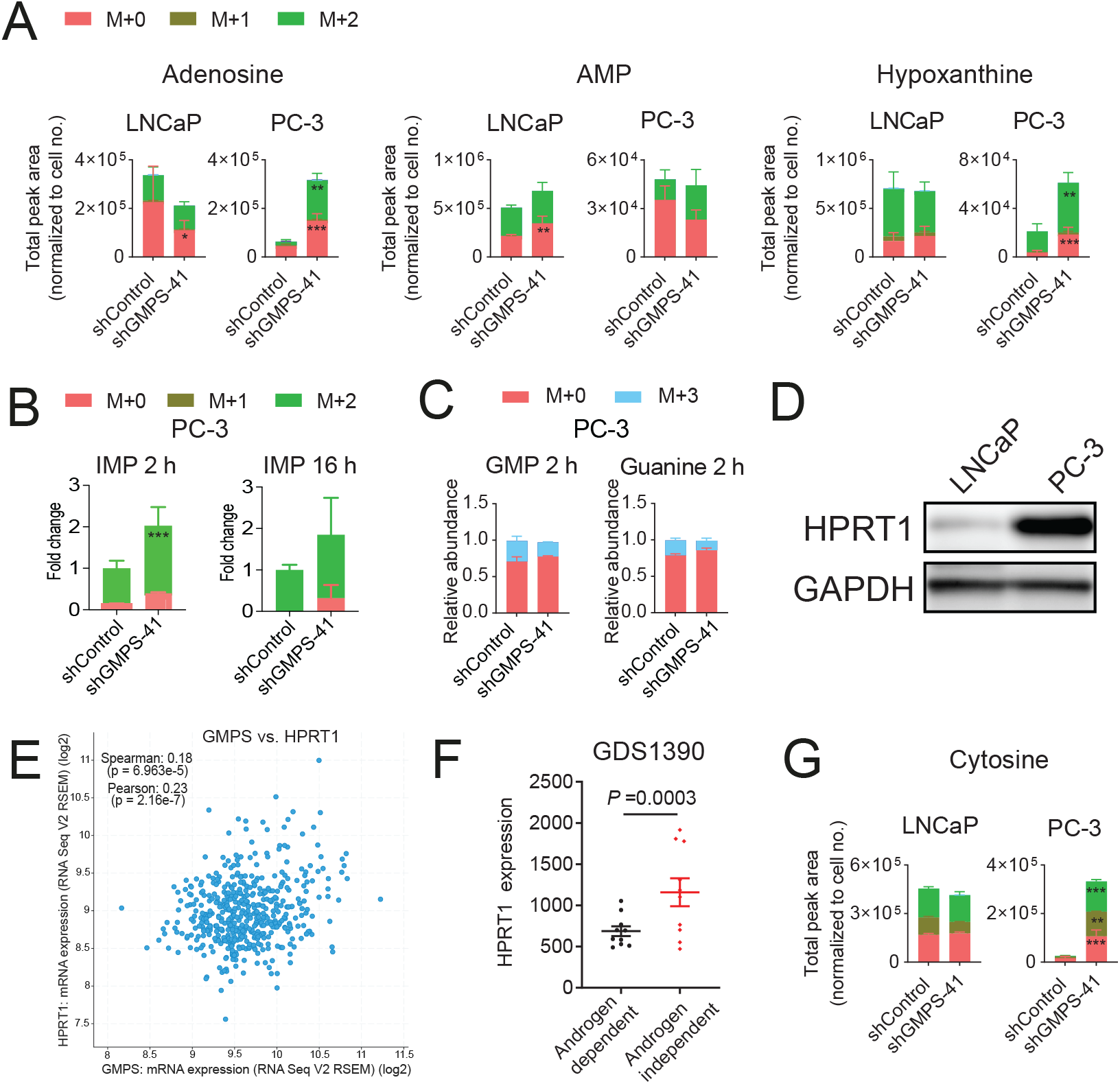
Abundance of cellular metabolites were measured using LC-MS in LNCaP and PC-3 shControl or shGMPS cells. (A) Analysis of ^15^N-(amide)-glutamine derived cellular purine metabolites adenosine, AMP and hypoxanthine. (B) ^15^N-(amide)-glutamine derived purine metabolite IMP in PC-3 cells after 2 h and 16 h incubation. Two-way ANOVA test.*, P < 0.05; **, P < 0.01; ***, P < 0.001. (C) Relative abundance of ^15^N-(amide)-glutamine derived GMP and guanine in PC-3 cells after 2 h and 16 h incubation. (D) HPRT1 protein was detected by western blotting in LNCaP and PC-3 cells. GAPDH was used as the loading control. (E) GMPS mRNA expression correlates with HPRT1 in TCGA dataset (cBioPortal). (F) HPRT1 mRNA expression in androgen dependent (n = 12) and androgen independent (n = 49) prostate cancer samples of GDS1390 dataset. Data are mean ± SEM; unpaired parametric t-test. (G), Analysis of ^15^N-(amide)-glutamine derived cellular pyrimidine metabolite cytosine. Two-way ANOVA test.*, P < 0.05; **, P < 0.01; ***, P < 0.001.

